# Ghrelin signaling regulates feeding behavior, metabolism, and memory through the vagus nerve

**DOI:** 10.1101/2020.06.16.155762

**Authors:** Elizabeth A. Davis, Hallie S. Wald, Andrea N. Suarez, Jasenka Zubcevic, Clarissa M. Liu, Alyssa M. Cortella, Anna K. Kamitakahara, Jaimie W. Polson, Myrtha Arnold, Harvey J. Grill, Guillaume de Lartigue, Scott E. Kanoski

## Abstract

Vagal afferent neuron (VAN) signaling sends information from the gut to the brain and is fundamental in the neural control of feeding behavior and metabolism. Recent findings reveal that VAN signaling also plays a critical role in cognitive processes, including hippocampus (HPC)-dependent memory. VANs, located in nodose ganglia, express receptors for various gut-derived endocrine signals, however, the function of these receptors with regards to feeding behavior, metabolism, and memory control is poorly understood. We hypothesized that VAN-mediated processes are influenced by ghrelin, a stomach-derived orexigenic hormone, via communication to its receptor (growth hormone secretagogue receptor [GHSR]) expressed on gut-innervating VANs. To examine this hypothesis, rats received nodose ganglia injections of an adeno-associated virus (AAV) expressing short hairpin RNAs targeting GHSR (or a control AAV) for RNA interference-mediated VAN-specific GHSR knockdown. Results reveal that VAN GHSR knockdown induced various feeding and metabolic disturbances, including increased meal frequency, impaired glucose tolerance, delayed gastric emptying, and increased body weight compared to controls. Additionally, VAN-specific GHSR knockdown impaired HPC-dependent episodic contextual memory and reduced HPC brain-derived neurotrophic factor expression, but did not affect anxiety-like behavior or levels of general activity. A functional role for endogenous VAN GHSR signaling was further confirmed by results revealing that VAN signaling is required for the hyperphagic effects of ghrelin administered at dark onset, and that gut-restricted ghrelin-induced increases in VAN firing rate require intact VAN GHSR expression. Collective results reveal that VAN GHSR signaling is required for both normal feeding and metabolic function as well as HPC-dependent memory.

## INTRODUCTION

The vagus nerve is a primary conduit of communication between the gastrointestinal (GI) tract and the brain. Vagal afferent neurons (VANs) and their ascending sensory fibers of the vagus transmit information about gastric distension, calorie content, and peptide hormone release to the brain to regulate feeding behavior and metabolism [1]. VAN-mediated transmission of nutrient, metabolic, and physiological signals to the brain are hypothesized to occur, in part, via paracrine signaling from GI-derived peptides to receptors expressed on VAN terminals innervating the GI tract [2]. However, technical limitations in targeting VAN GI peptide receptors either pharmacologically or with transgenic models have precluded advancement in understanding the physiological role of VAN-mediated paracrine signaling. For example, receptors for the orexigenic gut hormone ghrelin (growth hormone secretagogue receptor; GHSR), which is released from the stomach epithelium in response to energy restriction and in anticipation of a meal [3-6], are expressed on VANs [7-10], including those that directly innervate the stomach [11, 12]. GHSRs are also dispersed throughout the brain [13, 14]. Based on this receptor localization profile, ghrelin has been purported to act through both a VAN-mediated paracrine as well as a blood circulation-to-brain (endocrine) pathway [7, 15]. However, the specific role of VAN GHSR paracrine signaling in regulating food intake and metabolic function is poorly understood.

Recent studies reveal that in addition to regulating feeding behavior and metabolic function, GI-derived VAN signaling also affects higher-order neurocognitive processes such as affective motivational behaviors and learning and memory systems [16-19]. For example, we recently revealed that selective ablation of GI-innervating VANs impaired memory processes that rely on the integrity of the HPC [17]. Given that ghrelin promotes memory processes via incompletely understood mechanisms [20-24], it is likely that VAN-mediated regulation of HPC function involves VAN ghrelin signaling. Here, we employ multiple levels of analysis to investigate the hypothesis that endogenous VAN ghrelin signaling regulates feeding behavior, metabolic processes, and promotes HPC-dependent memory function.

## RESULTS

### Ghrelin receptor mRNA (*Ghsr*) is expressed at similar levels in left and right nodose ganglion in both the rat and the mouse

Both rats and mice express *Ghsr* mRNA in the nodose ganglia [7-10], thus suggesting that the molecular machinery is present for ghrelin to bind to vagal afferent nerves to influence physiological processes. Here, we show that *Ghsr* expression did not differ between the left and the right nodose ganglia in either the rat (Fig. 1A) or in the mouse (Fig. 1B). While all subsequent results are from a rat model, we included data from both rats and mice in this experiment given a recent report that left vs. right VAN signaling differ with regards to neurocognitive outcomes and downstream neuroanatomical pathways in mice [19].

**Figure 1.**
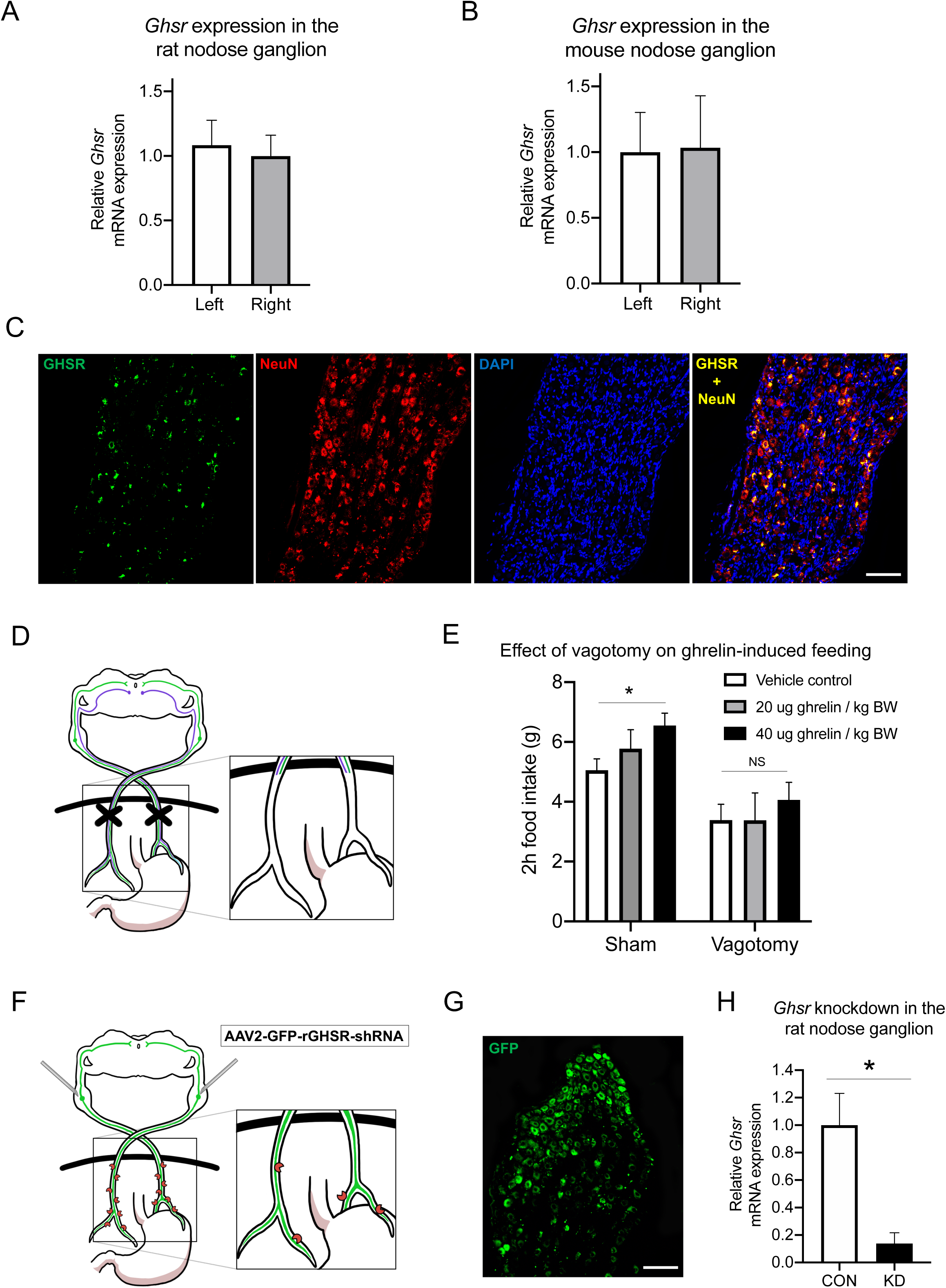
VAN signaling is required for the hyperphagic effects of peripheral ghrelin, and the ghrelin receptor (GHSR) is expressed in the nodose ganglion and is reduced with targeted RNA interference. The left and right nodose ganglia (LNG, RNG), which house VAN cell bodies, express *Ghsr* mRNA in both rats (A) and mice (B) with no differences in expression between LNG and RNG in either species. *Ghsr* mRNA (green) is exclusively expressed in rat nodose ganglion neurons (red, neuronal marker *NeuN*), as demonstrated co-localization of *Ghsr* and *NeuN* (yellow), and approximately ∼73% of *NeuN*+ nodose ganglion neurons co-express *Ghsr* (C). After complete bilateral subdiaphragmatic vagotomy in rats (D), the hyperphagic effect of IP ghrelin at 40μg/kg BW injected in the early dark cycle was abolished compared with controls (E). VAN *Ghsr* expression is knocked down in rats via bilateral nodose ganglia injections of a custom-designed AAV (AAV-2 GFP-rGHSR-shRNA) (F). Representative histology of VAN cell bodies in the nodose ganglion expressing the GHSR shRNA AAV with a green fluorescent protein (GFP) transgene (G). Gene expression analyses confirmed a statistically significant ∼86% knockdown of the *Ghsr* gene in the nodose ganglion (H). All data presented as mean +/- SEM.

### *Ghsr* is expressed in neurons but not glia in the rat nodose ganglion

Fluorescent *in situ* hybridization analyses reveal that *Ghsr* mRNA is expressed in neurons in both the left and the right rat nodose ganglion, as evident from colocalization of *Ghsr* with the neuronal marker *NeuN* (representative image in Fig. 1C). Quantitative analyses confirm that *Ghsr* is co-expressed in 73% of *NeuN*+ nodose ganglia neurons (SEM +/-7.6%). Further, consistent with data from humans [9], *Ghsr* is exclusively expressed in neurons and not glia in the rat nodose ganglion, as indicated by a total lack of *Ghsr* that is not colocalized within *NeuN*-expressing neurons.

### Vagus nerve signaling is required for the hyperphagic effects of peripheral ghrelin during the nocturnal cycle

We investigated whether the vagus nerve is functionally required for the peripheral orexigenic effects of ghrelin in the rat by comparing 2h cumulative nocturnal/dark cycle food intake following IP ghrelin injections (20μg/kg body weight (BW), 40μg/kg BW, or a saline/vehicle control) in rats that had a bilateral subdiaphragmatic vagotomy (SDV; Fig. 1D) compared with sham surgery. ANOVA results revealed that there was a main effect of surgery (Fig. 1E; p<0.05), and subsequent post hoc analyses revealed that the 40μg/kg dose significantly increased food intake relative to saline in the sham surgery group, but not in the SDV group (Fig. 1E). The 20μg/kg BW dose did not reach statistical significance for either surgical condition. These results indicate that the vagus nerve is required for the orexigenic effects of ghrelin when tested during the natural rodent feeding cycle (onset of dark cycle). SDV animals weighed significantly less than sham animals at the start of the IP ghrelin experiment (Sham: 409.8g +/- 8.5; SDV: 362.6g +/- 9.5; p<0.05) even after an extended post-surgical recovery period, as we have previously shown [25]. However, while SDV animals ate significantly less overall compared with the sham animals (Fig. 1E; p<0.05), food intake between groups in the saline condition was not significantly different when normalized per 100g body weight (p>0.05; Supp. Fig. 2A), suggesting that the lack of a ghrelin hyperphagic effect in the SDV group is unlikely to be based on a ceiling effect for maximal food intake.

### Nodose ganglion *Ghsr* expression is reduced with targeted RNA interference without affecting expression of other feeding-relevant peptide receptors

Based on the above molecular and behavioral results suggesting a physiological role for VAN ghrelin/GHSR signaling in feeding behavior, we developed an approach to knockdown GHSR specifically within VAN in the rat. Nodose ganglion histology analyses revealed that VAN cell bodies expressed the green fluorescent protein (GFP) transgene driven by the AAV (Fig. 1G). Bilateral nodose ganglion injections of a custom-designed AAV for *Ghsr*-targeted mRNA interference (AAV-2 GFP-rGHSR-shRNA) (Fig. 1F) were administered in rats, which significantly reduced VAN *Ghsr* mRNA expression by ∼86% compared with controls (Fig. 1H; p<0.05). To evaluate potential compensatory changes in expression of other feeding-relevant VAN receptors that are known to interact with GHSR [9], we evaluated melanin-concentrating hormone receptor 1 (*Mch1r*) and cannabinoid 1 receptor (*Cb1r*) mRNA expression in the nodose ganglia of rats that received *Ghsr* shRNA vs. controls. Results reveal no differences in *Mch1r* or *Cb1r* expression between GHSR knockdown and controls, which supports the selectivity of the VAN shRNA GHSR approach and further indicates that behavioral and metabolic effects of VAN GHSR knockdown are unlikely to be secondary to chronic changes in VAN *Mch1r* or *Cb1r* signaling.

### VAN-specific GHSR knockdown blocks the VAN neural response to gut-restricted ghrelin

To investigate whether ghrelin administration restricted to the GI tract evokes VAN neural response via GHSR, a modified *in situ* model of a decorticated artificially perfused rat (DAPR; Fig. 2A) was used. The modification allowed for separate perfusion above and below the diaphragm. Using this approach, we were able to simultaneously record both left and right nodose ganglia neural activity in response to an intra-arterial injection of ghrelin (5nmol vs. saline control) restricted to the GI tract. Representative traces show responses to ghrelin from an animal with the GHSR knockdown in the right nodose ganglion (RNG) with control AAV in the left nodose ganglion (LNG) (Fig. 2B), as well as from an animal with the GHSR knockdown in the LNG and the RNG as a control (Fig. 2C). When maximum response to ghrelin were compared between vagal recordings of the control injected side, there were no differences between left and right responses (Fig. 2D). However, when comparing the vagal response between the GHSR knockdown and the contralateral side within-animal, there was a statistically significant decrease in maximum response on the knockdown side (Fig. 2E; p<0.05). Vagal afferent activity in response to intra-arterial injection of cholecystokinin (CCK; 1μg) within the same animals was also measured. Representative traces include responses to CCK from an animal with the GHSR knockdown in the RNG (LNG control) (Fig. 2F), and an animal with the GHSR knockdown in the LNG (RNG as control) (Fig. 2G). CCK infusion yielded a pronounced increase in VAN neural activity in both control and knockdown sides, with trend towards a greater maximum response to CCK in the right compared to left vagus nerve (p=0.09; Fig. 2H). However, there were no differences in maximum response to CCK between knockdown and controls (Fig. 2I). Collectively, these results support a paracrine ghrelin-vagal pathway by revealing that VAN respond to GI-restricted ghrelin in a GHSR-dependent manner, whereas VAN response to CCK is GHSR-independent. Given that ghrelin circulation in this experiment was precluded from supradiaphragmatic access to GHSRs expressed in the nodose ganglia, these results highlight a functional role for GHSRs expressed on subdiaphragmatic vagal afferent terminals innervating the GI tract.

**Figure 2.**
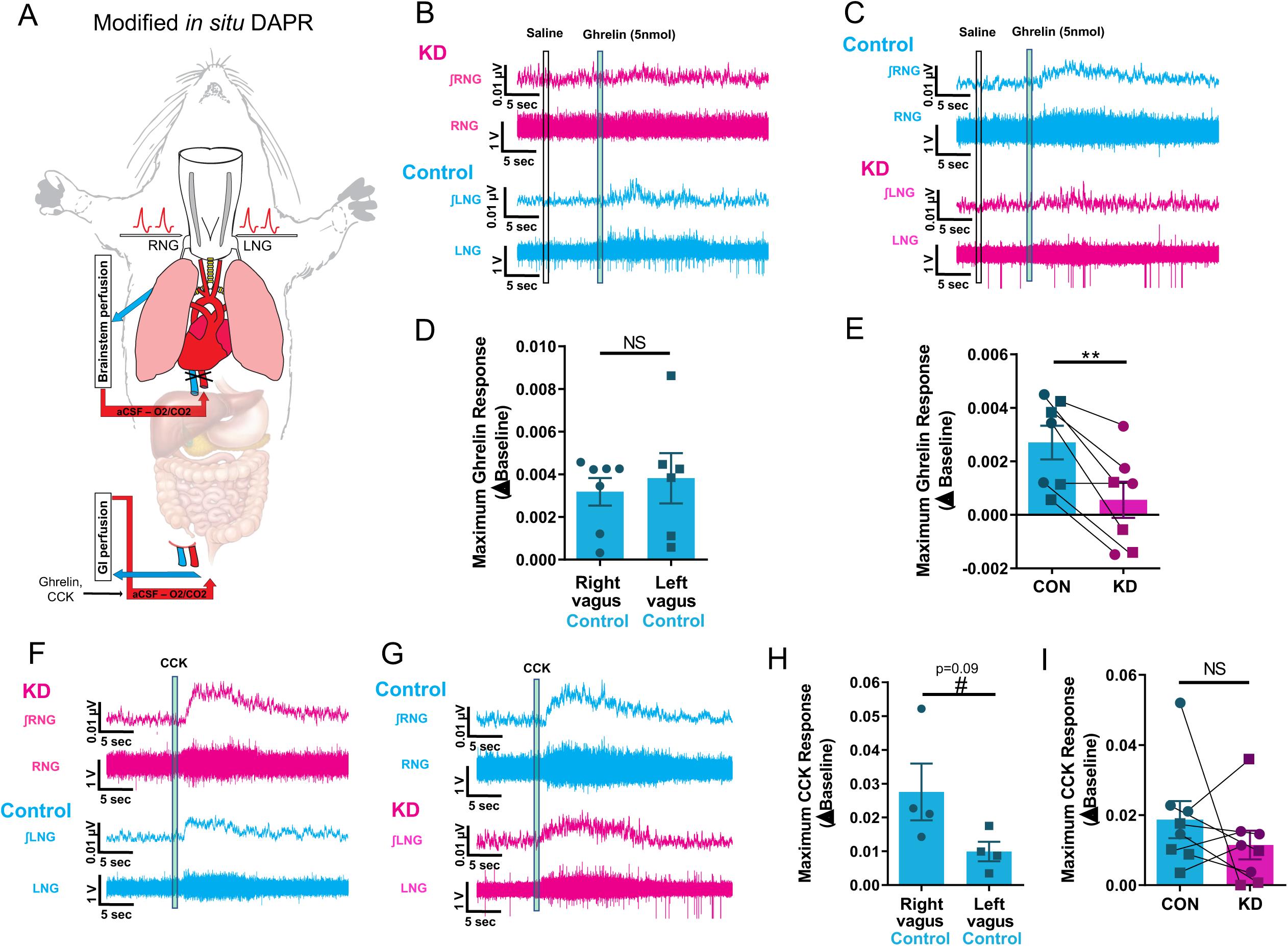
VAN-specific GHSR knockdown blocks the VAN neural response to gut-restricted ghrelin. An *in situ* model of a decorticated artificially perfused rat (DAPR) was used to simultaneously record both left and right supranodose afferent activity in response to exogenous infusions of saline, ghrelin or CCK restricted to the gastrointestinaI circulation (A). Representative traces of right (top) and left (bottom) vagal afferent activity within the same animal in response to intra-arterial injection of ghrelin (5nmol) (an animal with shRNA for GHSR in the RNG in B; an animal with shRNA for GHSR in the LNG in C). In the control AAV vagal side (cyan), ghrelin produces a small but reproducible and significant increase in vagal afferent activity, which is not present in the GHSR shRNA vagal side (magenta) (B and C). There is no difference in maximum ghrelin response between control injected left or right vagal recordings (D). Conditional knockdown of ghrelin receptor significantly blunts ghrelin induced vagal activity compared with control (E). Representative traces of right (top) and left (bottom) vagal afferent activity in response to intra-arterial injection of CCK (1μg) within the same animal (an animal with shRNA for GHSR in the RNG in F; an animal with shRNA for GHSR in the LNG in G). CCK produces a pronounced increase in vagal afferent activity in both control and shRNA injected sides (F and G). H) There is trend towards a greater maximum CCK response in the right compared to left vagus nerve (n=4, p=0.09). Knockdown of ghrelin receptor has no effect on CCK induced vagal activity compared to control (I).

### VAN-specific GHSR knockdown alters meal pattern but not cumulative 24h food intake

Meal-pattern analyses were conducted using automated food intake monitors to examine the effect of VAN-specific GHSR knockdown on meal size and meal frequency. Separate two-way repeated-measures ANOVAs for each dependent variable over 7 days (as well as an unpaired two-sample t-test for the 7-day averages of each dependent variable) revealed a significant increase in meal frequency in the VAN-specific GHSR knockdown group compared with controls (Figs. 3A and 3B; p<0.05), coupled with a nonsignificant trend in reduced average meal size (Figs. 3C and 3D; p=0.09) such that no significant differences were observed in cumulative 24h chow intake over 7-day investigated time period (Figs. 3E and 3F). VAN-specific GHSR knockdown had no significant effect on meal size, frequency, or cumulative intake when analyses were restricted to the light cycle (data not shown).

**Figure 3.**
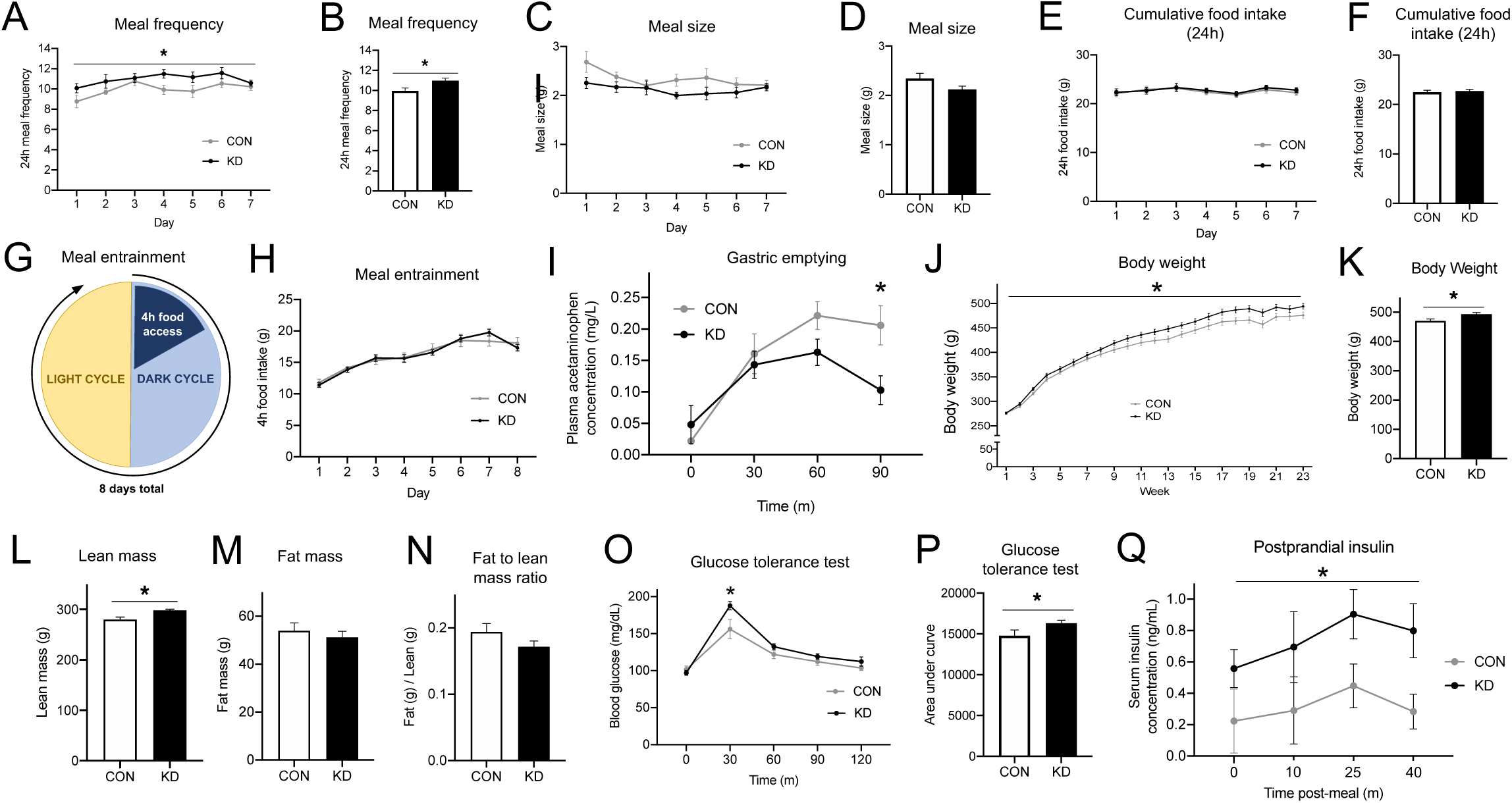
VAN-specific GHSR knockdown increases meal frequency, body weight and lean mass while disrupting gastric emptying rate and peripheral glucose tolerance. Compared with control animals, VAN-specific GHSR knockdown animals displayed an increased meal frequency (A, B) and a non-significant trend toward decreased meal size (C, D), such that there were no changes in 24h cumulative food intake (E, F). Under meal entrainment conditions (G), there were no significant differences in food intake between knockdown and control animals (H). Rate of gastric emptying was significantly decreased in knockdown animals compared with controls 90 minutes after meal consumption (I). VAN-specific GHSR knockdown increased body weight compared with controls over time (J) and at 23 weeks post-surgery (K), at which point body composition was evaluated. Results showed increased lean mass in knockdown animals (L). However, there were no differences between groups in fat mass (M) or fat to lean mass ratio (N). VAN-specific GHSR knockdown impaired glucose tolerance compared with controls in an IP glucose tolerance test at the 30-minute timepoint (O), and in total as measured by area under the curve (P). VAN-specific GHSR knockdown postprandial increased serum insulin levels compared with controls (Q). All data presented as mean +/- SEM.

### VAN-specific GHSR knockdown does not impair meal entrainment

Next, we examined the effect of VAN-specific GHSR knockdown on meal entrainment (diagram of feeding protocol in Fig. 3G). We restricted animals’ food access to a 4h period per day over 8 days, which requires animals to learn to consume all of their daily calories during a short period of time. Results showed no differences between VAN-specific GHSR knockdown and controls in total food consumed by day over the 8-day period (Fig. 3H; p<0.05). These findings suggest that VAN-specific GHSR knockdown impairs neither the learning required to adapt to meal entrainment. Furthermore, food intake data from Day 1 (prior to learning) reveal that there were no group differences following either 1h (Control = 5.62g [SEM 0.51], VAN GHSR KD = 5.45g [SEM 0.37]) or 4hrs (Fig. 3G) of refeeding after an unanticipated fast.

### VAN-specific GHSR knockdown slows gastric emptying rate

Using a gavage-based acetaminophen approach [26], we investigated the effect of VAN-specific GHSR knockdown on gastric emptying rate. A two-way repeated-measures ANOVA revealed a significant interaction between treatment and time, as well as a main effect of time (Fig. 3I; p<0.05). Post hoc analysis revealed a significant separation in means at the 90 min time point, indicating that VAN-specific GHSR knockdown slows gastric emptying compared with controls.

### VAN-specific GHSR knockdown increases body weight and lean mass

VAN-specific GHSR knockdown increased body weight over time (Fig. 3J; p<0.05) compared with controls. We therefore were interested in evaluating body composition, which was performed via nuclear magnetic resonance imaging at 23 weeks post-surgery. At the time of body composition evaluation, the knockdown animals weighed significantly more than controls (Fig. 3K; p<0.05), and had increased lean mass compared with controls (Fig. 3L; p<0.05). However, there were no differences in fat mass (Fig. 3M) nor the ratio of fat mass to lean mass (Fig. 3N), calculated by fat(g)/lean(g). Taken together, these results suggest that VAN-specific GHSR knockdown increases body weight, driven by increases in lean mass.

### VAN-specific GHSR knockdown impairs glucose tolerance and increases postprandial circulating insulin levels

We tested the effect of VAN-specific GHSR knockdown on intraperitoneal (IP) glucose tolerance. A two-way, repeated measures ANOVA (treatment x time) revealed a significant interaction between treatment and time, as well as a main effect of time. Post hoc analysis revealed a significant separation in means at the 30-minute time point (Fig. 3O; p<0.05), where knockdown animals had elevated blood glucose levels compared with controls. Analysis using the standard area under the curve method (which is traditionally used for glucose tolerance testing) similarly revealed that knockdown animals had increased blood glucose in response to IP glucose (Fig. 3P; p<0.05). These results indicate that VAN-specific GHSR knockdown impairs IP glucose tolerance compared with controls. To investigate potential underlying mechanism of the impaired glucose tolerance, we tested the effect of VAN-specific GHSR knockdown on postprandial serum insulin levels. A two-way, repeated measures ANOVA (treatment x time) revealed a significant main effect of group; knockdown animals had increased insulin levels compared with controls (Fig. 3Q; p<0.05). These results suggest that the impairment in glucose tolerance in VAN-specific GHSR knockdown animals is mediated by insulin intolerance and not by reduced pancreatic insulin production.

### VAN-specific GHSR knockdown impairs HPC-dependent contextual episodic memory and HPC neurotrophin levels without affecting anxiety-like behavior

Novel object in context (NOIC) is a rodent memory test of HPC-dependent contextual episodic memory (Fig. 4A). Results reveal that VAN-specific GHSR knockdown animals had a significantly reduced shift from baseline of novel object investigation relative to control animals (unpaired Student’s t-test; Fig. 4B; p<0.05). There was no difference between groups in the exploration of the non-novel object on test day (control: 18.51s +/- 0.96, knockdown: 21.38s+/- 1.77). These findings indicate that VAN GHSR signaling is required for HPC-dependent contextual episodic memory in rats.

**Figure 4.**
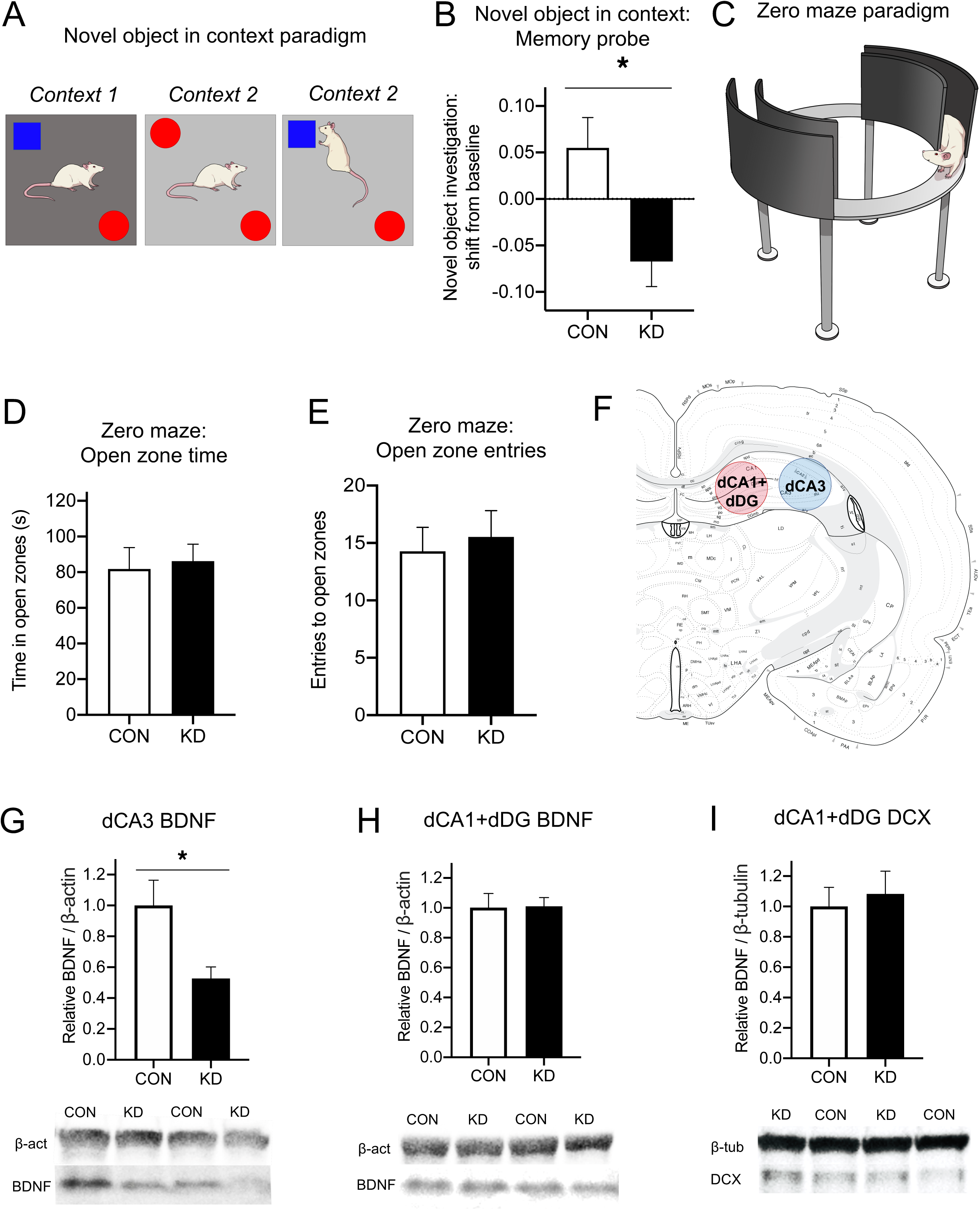
VAN-specific GHSR knockdown impairs HPC-dependent contextual episodic memory and reduces HPC CA3 BDNF without affecting anxiety-like behavior. In the novel object in context (NOIC) task (A), VAN-specific GHSR knockdown animals were significantly impaired in novel object exploration shift from baseline during the memory probe compared with controls (B), indicating that VAN-specific GHSR knockdown impairs HPC-dependent contextual episodic memory. In the zero maze task testing anxiety-like behavior (C), VAN-specific GHSR knockdown did not alter zero maze performance compared to controls, as measured by time in the open zones (D) and number of entries to open zones (E). Dorsal CA3 (dCA3) and dorsal CA1 plus and dorsal dentate gyrus (dCA1+dDG) lysates (F), were analyzed for protein levels associated with neuroplasticity and neurogenesis. The neuroplasticity-associated protein BDNF was decreased in knockdown animals compared with controls in dCA3 lysates (H), but there were no differences between groups in dCA1+dDG lysates. There were no differences in the proliferation-associated protein DCX between groups in the dCA1+dDG lysate (I), where the proliferating dentate gyrus neurons are located.

To confirm that decreased novel object exploration in VAN-specific GHSR knockdown animals during NOIC testing is not a general avoidance of novel objects due to altered anxiety, we tested anxiety-like behavior using the zero maze test (Fig. 4C). Results show no differences between VAN-specific GHSR knockdown animals and controls in anxiety-like behavior for both time in open zones (Fig. 4D) as well as open zone entries (Fig. 4E).

The HPC-dependent memory impairments seen in the NOIC test may be based, in part, on reduced neurotrophic signaling in the dorsal CA3 subregion. Immunoblot analyses of HPC subregions (Fig. 4F) revealed decreased protein levels of brain-derived neurotrophic factor (BDNF) in CA3-enriched brain tissue punches of VAN-specific GHSR knockdown animals compared with controls (unpaired Student’s t-test; Fig. 4G; p<0.05). No group differences were identified in brain punches enriched with the dCA1+dDG subregions for expression either of BDNF (Fig. 4H) or the proliferation marker, doublecortin (DCX; Fig. 4I).

### VAN-specific GHSR knockdown does not affect appetitive contextual or spatial memory

We investigated the effect of VAN-specific GHSR knockdown on a novel spatial foraging task (Supp. Fig. 1A), which tests the ability for animals to learn and remember the spatial location of palatable food. Results revealed no groups differences in latency to correct hole (Supp. Fig. 1B) nor errors before correct hole (Supp. Fig. 1C) during the training. There were also no group differences in retention during the memory probe, as measured by the correct + adjacent holes investigated / total holes investigated (Supp. Fig. 1D). In a different task testing conditioned place preference for a high fat diet (Supp. Fig. 1E), there were no differences between VAN-specific GHSR knockdown and controls in preference for food-paired context (time spent in context, shift from baseline) (Supp. Fig 1E).

## DISCUSSION

Paracrine vagus nerve signaling by gut hormones is a critical pathway through which the GI tract communicates to the brain to regulate energy balance and metabolic function [27, 28]. However, little is understood about the neurobiological mechanisms and physiological relevance of VAN paracrine signaling. Recent reports identify a physiological role for leptin and GLP-1 in the control of feeding behavior and metabolic outcomes via VAN signaling [29, 30], however, whether the stomach-derived orexigenic hormone ghrelin signals through a vagal paracrine pathway is poorly understood. The present results reveal that VAN GHSR signaling regulates various aspects of feeding behavior and metabolic processes. VAN-specific GHSR knockdown increased meal frequency with a trend toward decreased meal size compared with controls such that there were no differences in cumulative 24h food intake between groups. Additionally, metabolic parameters were disrupted in knockdown animals compared to controls, including impaired glucose tolerance driven by insulin resistance, slower gastric emptying rate, and increased body weight driven by increased lean mass. Overall, these results suggest that VAN ghrelin/GHSR signaling plays an endogenous role in normal feeding behavior, metabolism, and memory.

VAN-specific GHSR knockdown also impaired hippocampal-dependent contextual episodic memory. These findings provide a neurobiological mechanism for our previous results in which both complete subdiaphragmatic vagotomy and GI-specific vagal afferent ablation impair hippocampal function [17]. It is possible that the observed impairments in episodic memory are functionally related to the increased spontaneous meal frequency following VAN-specific GHSR knockdown. Lesioning or inhibition of hippocampal neurons reduces the intermeal interval in rodent models, in addition to increasing intake at the next meal [31-34]. In humans, interfering with meal-related episodic memory (by decreasing perceived food intake while controlling for actual food intake) increases subjects’ self-reported hunger rating during the intermeal interval [35], while a separate study showed that simply being asked to recall a previous meal decreases future intake at the subsequent meal [36]. However, it is also possible that the increased meal frequency effects are based, in part, on the reduced gastric emptying rate in GHSR VAN knockdown rats, which may contribute to reduced meal size with a compensatory increase in meal frequency. However, this possibility is less likely as the trend for reduced spontaneous meal size in the present study failed to reach statistical significance. These results identify VAN GHSR signaling as a potential critical physiological link between episodic memory and feeding behavior.

Based on our previous neuroanatomical pathway tracing and molecular results, it is probable that GHSR VAN signaling engages caudal mNTS neurons that project to the medial septum, which in turn project to the HPC CA3 (dCA3) subregion to influence neurotrophic signaling and memory function [17]. Consistent with this framework, the current study revealed decreased protein levels of the neurotrophic protein BDNF in the dCA3 HPC subregion (but not CA1 or DG subregions) following VAN-specific GHSR knockdown, which highlights a potential molecular mechanism for the contextual episodic memory impairments seen after VAN GHSR knockdown given the established role for HPC BDNF signaling in memory control [37, 38]. A potential driver for these changes in HPC BDNF is the increase in body weight observed after VAN-specific GHSR knockdown. Increased body weight is associated with reduced leptin receptor signaling in the brain [39], and leptin receptor signaling has been shown to increase central *Bdnf* mRNA [40]. However, this is unlikely as the body weight gain associated with VAN GHSR knockdown is modest compared to the obese state that is commonly associated with central leptin resistance.

Previous work shows that ghrelin blocks the downregulation of VAN *Cb1r* and *Mch1r* expression that occurs following refeeding after a fast [9]. Thus, it may be the case that the increased meal frequency associated with VAN-specific GHSR knockdown is based, in part, on direct interactions between VAN GHSR signaling and these feeding-relevant receptor systems. However, we did not find changes in *Mch1r* nor *Cb1r* gene expression in VAN GHSR KD rats compared with controls, suggesting that dysregulation of these receptors is not contributing to the meal pattern changes observed. Notably, ablation of cholecystokinin (CCK) receptor-expressing VAN, which overlap substantially with GHSR-positive VAN [9], produces similar increases in meal frequency without affecting meal size [41]. While CCKR expression was not measured in the present study, the possibility that GHSR and CCKR interact in VAN to influence meal frequency is unlikely given that VAN GHSR KD did not affect VAN neural responses to gut-restricted CCK.

In addition to the vagal paracrine pathway investigated in the present study, ghrelin also influences food intake, metabolism, and memory function via a putative blood-to-brain pathway, as well as through non-vagal peripheral signaling pathways [42-45]. Present results combined with previous research indicates that VAN and brain GHSR-mediated effects on feeding are divergent. For example, activation of GHSR signaling in the ventral hippocampus potently increases dark cycle food intake selectively via an increase in spontaneous meal size without influencing meal frequency [46], whereas VAN GHSR signaling in the present study predominantly increased meal frequency without affecting 24h cumulative food intake. In addition, present data show that VAN-specific GHSR knockdown does not influence food intake in a meal entrainment schedule that has been shown to increase peripheral ghrelin release immediately prior to meal access [6]. Food consumption in meal-entrained rats, however, is reduced by ventral hippocampus GHSR blockade [47], thus highlighting the importance of brain but not VAN GHSR signaling in ghrelin-mediated meal-entrained feeding. Overall, our results support emerging evidence that there are differential effects of ghrelin when acting through the VAN paracrine pathway versus the endocrine blood-to-brain pathway. However, these ventral hippocampus results were based on GHSR pharmacology. Due to the high endogenous constitutive activity of GHSR [48], the present findings may also be driven by changes in VAN activity typically driven by constitutive GHSR activity, which may, in part, contribute to the discrepancies between VAN and brain GHSR effects on feeding. Regardless, the present study supports a unique role for VAN ghrelin/GHSR signaling that is functionally distinct from ghrelin/GHSR signaling in the brain with regards to meal patterns and entrained feeding.

Electrophysiological results from the present study reveal that ghrelin functionally excites VAN neural responses in a GHSR-dependent fashion. Given that ghrelin circulation in this experiment was precluded from supradiaphragmatic access to GHSRs expressed in the nodose ganglia, these results are likely based on GHSRs expressed on subdiaphragmatic vagal afferent terminals innervating the GI tract. The approach used to obtain these results is novel, as to our knowledge this is the first instance of simultaneous recordings from both vagal nerves being performed, as well as the first application of VAN neural recording in response to a gut-restricted circulating hormone. Notably, we found that there were no laterality differences in ghrelin-induced neural responses between the left and right vagus nerve, which is consistent with our gene expression data showing no differences in *Ghsr* mRNA expression when comparing the left and right nodose ganglion in both mice and rats. We further revealed that VAN GHSR knockdown reduced the VAN neural response to ghrelin, but not CCK, thus further confirming the specificity of the GHSR-targeted RNA interference approach. Complementing present findings that ghrelin engages VAN signaling both electrophysiologically and functionally, our results show that SDV attenuates the hyperphagic effects of peripheral ghrelin during the early nocturnal phase. These findings are consistent with a previous report by Date *et al*. [7], which found that both SDV and chemical vagal deafferentation attenuate the hyperphagic effect of ghrelin. However, in the study by Date and colleagues the evaluation of ghrelin-stimulated hyperphagia occurred before the complete recovery from SDV surgery, which is critical for the return to feeding behavior and other physiological functions after vagotomy [25, 49-51]. In contrast, Arnold et al., reported that SDV has no effect on ghrelin-mediated hyperphagia in rats [52]. These discrepancies may be based on methodological differences between experiments, as the present study was performed in the dark cycle, whereas this latter study by Arnold and colleagues was performed in the light cycle. A potential effect of photoperiod may be, in part, due to the diurnal rhythm of nodose ganglion *Ghsr* mRNA expression, with expression higher during the light cycle compared with the dark cycle [10]. Differences in results between studies may also be based on circadian variation in gastric VAN mechanosensitivity, which is increased during the light cycle and decreased during the dark cycle [53]. Another important methodological difference between the present study and the study by Arnold *et al.* is that here we examined the orexigenic effects of ghrelin using laboratory chow, while Arnold and colleagues used a liquid diet. Future research will be required to determine the importance of diet composition and viscosity, photoperiod, and other experimental parameters on the role of the vagus nerve in mediating the orexigenic effect of exogenous ghrelin.

The present results collectively reveal a novel role for endogenous VAN GHSR signaling in multiple domains, including feeding behavior, metabolism, and HPC-dependent memory. These findings not only identify a novel neurobiological mechanism for ghrelin’s effects on various physiological and behavioral processes, but are also consistent with emerging evidence that gut peptides engage in functionally-relevant paracrine gut-to-brain signaling via the vagus nerve. Future research on the importance of VAN GHSR signaling and its translational potential is merited.

## ACKNOWLEDGEMENTS

This study was supported by the National Institute of Health grants: DK104897 (SK), DK118402 (SK), DK116558 (AS), DK118944 (CL), DK120162 (HW), DK021397 (HG), AT010192 (JZ), DK094871 (GL), DK116004 (GL), and the University of Sydney Special Studies Program (JP).

## AUTHOR CONTRIBTUTIONS

SK, GL, ED, HG, HW, and JZ designed experiments; GD and MA performed nodose injection procedures; ED, AN, and HW assisted with nodose injection procedures; AN performed vagotomies; JZ and JP performed DAPR; ED, HW, AN, and CL performed behavioral experiments; ED, AK, AS, HW, JZ, and JP harvested tissues; ED performed gene expression experiments; AC and ED performed immunoblotting experiments; AC performed FISH experiments; ED, JZ, GD, HW, and SK analyzed the data; ED drafted the manuscript; AC provided the figure art; SK edited the manuscript; all authors reviewed and approved of the manuscript.

## STAR METHODS

### Animals

For electrophysiological experiments, male Wistar rats (Charles River; 3-6 weeks old, 50-120g) were housed in specific-pathogen free cages on a 12h:12h light/dark cycle and had access to standard rat chow *ad libitum*. All procedures were approved by the University of Florida Institute of Animal Care and Use Committee.

For the gastric emptying experiment, adult male Sprague-Dawley rats (Charles River; 250-265g on arrival) were individually housed with *ad libitum* access (except where noted) to water and rodent chow (Purina 5001) on 12h/2h light/dark cycle. All procedures were approved by the University of Pennsylvania Institutional Animal Care and Use Committee.

For evaluation of *Ghsr* expression in the nodose ganglia of mice, adult male C57BL/6J mice (originally obtained from Jackson, bred in-house) were group housed with *ad libitum* access to rodent diet (PicoLab Rodent Diet 20, #5053) on a 13:11 hour light/dark cycle. All procedures were approved by the Children’s Hospital of Los Angeles Institute of Animal Care and Use Committee.

For all other experiments, Adult male Sprague–Dawley rats (Envigo; 250-275g on arrival) were individually housed with *ad libitum* access (except where noted) to water and chow (LabDiet 5001, LabDiet, St. Louis, MO) on 12h:12h light/dark cycle. All procedures were approved by the University of Southern California Institute of Animal Care and Use Committee.

### Vagotomy

Rats were habituated to liquid diet (Research Diets; AIN76A) for one day prior to surgery. Following a 24h fast and under ketamine (90mg/kg), xylazine (2.7mg/kg), and acepromazine (0.64mg/kg) anesthesia and analgesia (a subcutaneous injection of carprofen [5mg/kg]), complete subdiaphragmatic vagotomy (SDV) was performed (n=6) as described previously [54]. Body weight between groups was not significantly different at time of surgery (Sham: 310.9g +/- 3.2; SDV: 314.2g +/- 3.6). Briefly, a midline abdominal incision was made, the stomach was retracted caudally, and the liver was retracted cranially to expose the esophagus. The dorsal and ventral trunks of the vagus were then dissected from the esophagus. Each vagal trunk was ligated twice with a surgical thread at an interval of 1-2 cm, and then cauterized between the ligatures. In sham surgeries (n=7), the trunks were exposed but the vagus nerve was not ligated or cauterized. The incision was then closed with running sutures along the abdominal wall and stop sutures along the skin. Rats were allowed to recover on liquid diet for 3 days, and then were maintained on a powered chow diet for the remaining recovery period up to two weeks post-surgery, then were returned to a standard chow diet. Behavioral testing was performed approximately 3 months after surgery. After behavioral testing, SDV was verified functionally with intraperitoneal cholecystokinin (CCK)-induced food intake reduction as described [55, 56]. Briefly, the functional verification consists of analysis of food intake following intraperitoneal (IP) cholecystokinin (CCK-8, 2μg/kg; Bachem, Torrance, CA) or saline injections (treatments given counterbalanced on separate days) after an overnight fast. SDV rats were included in the statistical analysis if CCK treatment resulted in a less than 30% reduction of their food intake, as described [57]. No animals were removed for analyses based on these criteria.

### Food analyses following intraperitoneal ghrelin administration

The effect of vagotomy on IP ghrelin-mediated hyperphagia was measured in rats maintained on ad libitum chow and water. Food was removed 1h prior to injections. IP ghrelin (0, 20, or 40μg/kg body weight, Bachem) or saline (control) was administered immediately prior to dark onset. Food was returned at dark onset and cumulative 2h chow intake was recorded. Each animal received all three treatments (within-subjects design, counterbalanced order, vagotomy n=6, sham n=7) with each treatment day separated by 2 days.

### RNA interference-mediated VAN-specific GHSR knockdown

For *in vivo* knockdown of *Ghsr* gene expression, short hairpin RNA (shRNA) targeting *Ghsr* mRNA was cloned and packaged into an adeno-associated virus (AAV2; Vector Biolabs, Malvern, PA) and co-expressing green fluorescent protein (GFP), with both casettes downstream of the U6 promoter (titer 1⁄4 1.7e13 GC/mL; AAV2-GFP-U6-rGHSR-shRNA). The sequence of the shRNA is as follows: CCGGACTGCAACCTGGTGTCCTTTGCTCGAGCAAGGACACCAGGTTGCAGTTTTTG. A scrambled shRNA, GFP-expressing AAV2 downstream of a U6 promoter (titer 1⁄4 1.7e13 GC/mL; AAV2-GFP-U6-Scrmb-shRNA) was used as a control (Vector Biolabs, Malvern, PA).

Nodose ganglion injections were performed as previously described [17, 29]. Briefly, the day before surgery for nodose ganglia AAV injections, chow was removed and rats were given 130mL of diluted condensed milk. The day of surgery, rats were anesthetized via intramuscular injections of ketamine (90mg/kg), xylazine (2.8mg/kg), and acepromazine (0.72mg/kg) and were given a preoperative dose of analgesic (a subcutaneous injection of carprofen [5mg/kg]). Once the appropriate plane of anesthesia was reached, a midline incision was made along the length of the neck. The vagus nerve was separated from the carotid artery with Graefe forceps until the nodose ganglion was visible and accessible. A glass capillary (20μm-50μm tip, beveled 30° angle) attached to a micromanipulator was used to position and puncture the nodose ganglion and 1.5μL volume of AAV2-GFP-U6-rGHSR-shRNA (or the control AAV) was injected with a Picospritzer III injector (Parker Hannifin; Cleveland, OH) within the nodose ganglion at two sites: 0.75μL of virus was delivered rostral to the laryngeal nerve branch, and the remaining 0.75μL of virus was delivered caudal to the laryngeal nerve branch. The same procedure was performed for both nodose ganglia before the skin was closed with interrupted sutures. Postoperatively, rats were recovered to sternal recumbency on a heat pad and then returned to their home cage. Analgesic (5mg/mL carprofen subcutaneously) was given once per day for 3 days after surgery. Condensed milk was given on day 1 and day 2 postoperatively (130mL/day), with chow reintroduced on day 3 alongside the 130mL condensed milk. On day 4 and onwards, rats were returned to *ad libitum* access to chow. All subsequent analyses occurred after three weeks of post-surgical viral transfection. While our AAV was applied directly to the nodose ganglia, the subsequent RNA interference process would interfere with all cytoplasmic gene expression, including for the production GHSR protein that is trafficked to the vagal terminals. The DAPR electrophysiological approach (see next section) was used to further investigate the functional VAN response to ghrelin at the level of the terminals after GHSR knockdown.

### Modified decorticated artificially perfused rat (DAPR) preparation for gut-restricted ghrelin infusion and afferent vagal nerve recordings in situ

Wistar rats (male, 4 weeks old, 50-60g, N=8) received unilateral injection of AAV2-GFP-U6-rGHSR-shRNA in either the left (LNG) or right (RNG) nodose ganglia (n=4 per group) and a control virus (AAV-hSyn-ChR2(h134R)-eYFP) on the contralateral side. Three weeks after viral injections in nodose ganglia, we performed direct electrophysiological recordings of afferent vagal activity rostral to both LNG and RNG simultaneously using a modified decorticated artificially perfused rat (DAPR) *in situ* preparation 2 weeks following viral injections in nodose ganglia. The modification to the previously described DAPR [58-60] included retention of an intact GI tract with a separate perfusion system allowing specific targeting of GI vagal afferents by delivering ghrelin or CCK via arterial GI circulation (see schematics of the modified *in situ* DAPR in Fig. 2A). Wistar rats (male, 6 weeks old, 100-120g, n=8) were briefly anesthetized with isoflurane (4%), exsanguinated, decorticated (decerebrated at the precollicular level as originally described in [61]), and immediately submerged into ice-chilled artificial cerebrospinal fluid (aCSF; composition in mM: 125 NaCl, 24 NaHCO3, 3KCl, 2.5 CaCl2, 1.25 MgSO4 and 1.25 KH2PO4, 10 dextrose, pH 7.3) (Sigma-Aldrich and Fisher, USA). The skin was removed and the chest cavity was cut open keeping the diaphragm intact. Inferior vena cava and descending aorta were clamped immediately rostral to the diaphragm to isolate the brainstem from GI circulation (Fig. 2A). The descending aorta was cannulated immediately above the clamp with a double lumen catheter (Braintree Scientific) for rostral perfusion of the brainstem as previously described [60]. Below the diaphragm, descending aorta was cannulated with a separate double lumen catheter (Braintree Scientific) immediately above the iliac bifurcation for perfusion of the GI tract (Fig. 2A). The adjacent abdominal vein was severed in the most caudal lumbar region in order to allow release of the GI perfusate and its collection into a separate chamber for recirculation throughout the experiment. Both the right and left nerves were transected rostral to the ganglia to ensure recordings were of exclusively afferent vagal activity in both cervical vagal trunks simultaneously, without interference of the parasympathetic efferent signals that also travel via the vagus nerve. Following the surgery, the preparation was transferred to an acrylic perfusion chamber, and brainstem and GI were immediately perfused via the two implanted cannulae using separate warmed Ringer’s solutions (32-33°C) containing Ficoll PM70 (1.25%, Sigma-Aldrich, USA) bubbled with 95% O_2_/5% CO_2_ and pumped through in-line bubble traps and a filter (polypropylene mesh; pore size: 40 µm, Millipore) via two peristaltic pumps. Neuromuscular paralysis in the region above diaphragm was produced by addition of vecuronium bromide (2-4 µg/ml, Bedford Laboratories, USA) directly to the brainstem perfusate to eliminate the movement noise on nerve recordings. Perfusion pressures were measured via one lumen of the double-lumen catheters using two pressure transducers connected to an amplifier. Simultaneous recordings of the left and right afferent nerve activity (LNG and RNG, Fig, 2A) were obtained using glass suction electrodes (tip diameter, 0.2–0.3 mm), amplified (20–50K) and filtered (3-30K), sampled at 5 kHz (CED, Cambridge) and monitored using Spike2 (CED). The perfusate flows (19-24 ml/min for brainstem and 1.5-3.5 ml/min for GI perfusion) were adjusted to mimic physiologic perfusion levels and maintain the health of the preparation [58-60, 62]. Vasopressin (1.25-4 nM final concentration, Sigma-Aldrich, USA) was added to the brainstem perfusate to increase vascular resistance and aid in maintenance of adequate brainstem perfusion to preserve autonomic reflexes [58-60]. Bilateral vagal afferent responses were simultaneously recorded in response to slow bolus intra-arterial GI infusion of 1) saline (0.9%, 100μl), 2) ghrelin (Phoenix Peptide, 031-31, 5nmol in 100µl saline), and 3) CCK (Bachem BioScience, 4033010, 1μg in 100μl saline) within each preparation, always in this order. To account for the 400μl dead space in the tubing, 400μl saline was additionally administered after each infusion. CCK was delivered 5 min following the ghrelin injection to assess specificity of shRNA mediated knockdown.

For data analysis and comparison, the maximum amplitude of immediate afferent vagal nerve activation of integrated ∫LNG and ∫RNG (time constant=50 ms) to ghrelin or CCK administration in the GI tract was calculated as change (Δ) in μV from baseline representing the period of nerve activity recorded immediately following control saline injection. Comparisons were made either 1) between shRNA and control injected nodose ganglia within every animal, or 2) between left and right control injected nodose ganglia between subjects, and averaged for all preparations. Immediately following recordings, nodose ganglia were harvested for quantification of knockdown via gene expression (following three weeks of post-surgical viral transfection) (control n=12 ganglia, knockdown n=8 ganglia, left/right counterbalanced).

### Meal pattern

Meal size, meal frequency, and cumulative 24h food intake was measured using the Biodaq automated food intake monitors (Research Diets, New Brunswick, NJ). Animals (control n=13, knockdown n=12) were acclimated to the Biodaq on ad libitum chow for 3 days. Data were collected over a 7-day period (meal parameters: minimum meal size=0.2g, maximum intermeal interval=600s).

### Meal entrainment

Animals were subjected to a meal-entrainment schedule as previously described [47]. Briefly, animals (knockdown n=13, control n=11) were limited to chow access for 4h daily (for the first 4 hours of the dark cycle) over an 8-day period. Cumulative 4h food intake was recorded daily and water was available *ad libitum*.

### Gastric emptying

A gavage-based acetaminophen approach was used to measure gastric emptying rate, as this approach is standard in field for both humans and rats [26, 63, 64]. Animals (knockdown n=6, control n=6) were habituated to gavage prior to testing. Food was removed 16h before testing (last 4h of dark cycle plus full 12h light cycle) to limit the influence of variability in stomach contents on gastric emptying rate. At dark onset, rats were gavaged with 6 mL of a test meal of vanilla-flavored Ensure (Abbott Laboratories, Chicago, IL; 1.42 kcal/ml) containing 40 mg acetaminophen (Sigma-Aldrich, Cat #PHR1005). Tail vein blood (∼200µl) was collected immediately prior to dark onset/gavage (0; baseline) and 30, 60, and 90 min after dark onset/gavage with pre-coated EDTA microvette (Sarstedt, Nümbrecht Germany, supplier Fisher Scientific Cat# NC9976871). Tubes were immediately placed on wet ice, centrifuged, and plasma was collected and stored at −80°C until further processing. Acetaminophen concentrations were measured with a commercial kit (Cambridge Life Sciences, Ely, England K8003) adapted for a multi-well plate reader (Tecan Sunrise, Männedorf, Switzerland) according to manufacturer’s instructions. Each sample was run in duplicate.

### Body weight and body composition

Throughout all knockdown experiments, animals (knockdown n=13, control n=11) were weighed daily just prior to dark cycle onset to examine the effect of VAN-specific GHSR knockdown on body weight. At the conclusion of behavioral experiments, the Bruker LF90II nuclear magnetic resonance (NMR) minispec was used for non-invasive measurement of fat mass and lean mass to determine the effect of VAN-specific GHSR knockdown on body composition.

### Glucose tolerance test

Animals (knockdown n=12, control n=11) were food restricted 20 hours prior to an intraperitoneal glucose tolerance test (IP-GTT). Immediately prior to the test, baseline blood glucose readings were obtained from the tail tip and recorded by a blood glucose meter (One Touch Ultra2, LifeScan, Inc., Milpitas, CA). Each animal was then injected intraperitoneally (IP) with dextrose solution (1g dextrose / kg BW). Blood glucose readings were obtained at 30, 60, 90, and 120 min after IP injections.

### Postprandial serum insulin

Animals (knockdown n=8, control n=6) were food restricted 24 hours prior to a postprandial serum insulin test. Immediately prior to the test (at dark onset), baseline blood collections were taken from the tip of the tail (timepoint 0). Each animal was then allowed to consume the entirety of a 3g meal of powdered rodent chow, to which they had previously been habituated. All animals finished the meal between 15 and 20 min post meal access. Blood samples were then obtained from the tip of the tail at 10, 25, and 40 min after meal termination. Blood samples were allowed to clot at room temperature, centrifuged, serum was collected and stored at −80°C until further processing. Serum insulin concentrations were measured with a commercial kit (Crystal Chem Inc., Elk Grove Village, IL; Cat #90010) adapted for a multi-well plate reader (BioRad iMark Microplate Reader, BioRad Hercules, CA) according to manufacturer’s instructions. Each sample was run in duplicate.

### Novel object in context

To test the effect of VAN-specific GHSR knockdown on HPC-dependent contextual episodic memory, animals (knockdown n=10, control n=11) were tested in the Novel Object in Context (NOIC) procedure. Briefly, rats are habituated to Context 1, a semi-transparent box (15 in W × 24 in L × 12 in H) with orange stripes and Context 2, a gray opaque box (17 in W × 17 in L × 16 in H). Contexts and objects are cleaned with 10% ETOH between each animal. On Day 1following habituation (D1) at dark onset prior to the first meal, each animal is exposed to two distinct objects: a Coca Cola can (Object A) and a stemless wine glass (Object B) for 5 min in Context 1. On Day 2, the animals are exposed to duplicates of either Object A or Object B (counterbalanced by experimental group) for 5 min in Context 2. On Day 3 (D3), the animals are placed in the previous day’s location (Context 2) with Object A and Object B during a 5 min test period. Investigation time (*T*_*I*_) of both objects is measured by AnyMaze Behavior Tracking Software (Stoelting Co., Wood Dale, IL). Investigation is defined as sniffing or touching the object with the nose or forepaws. The task is scored by calculating the novel object investigation shift from baseline (from D1). If Object A is the novel object, shift from baseline is calculated as:

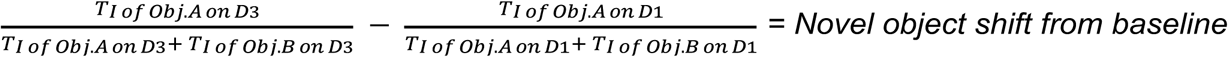

Normal rats will preferentially investigate the object that had not been previously seen in Context 2, given that it is a familiar object that is now presented in a novel context. This will result in a novel object shift from baseline that is increased compared with zero.

### Zero maze

To examine the effect of VAN-specific GHSR knockdown on anxiety-like behavior, animals (knockdown n=13, control n=11) were tested in the zero maze task. The zero maze apparatus is an elevated circular track, divided into four equal length sections. Two zones are open with 3 cm high curbs (‘open zones’), whereas the two other zones are closed with 17.5 cm high walls (‘closed zones’). Animals were placed in the maze for a single, 5 min trial, in which animal location was measured by AnyMaze Behavior Tracking Software (Stoelting Co., Wood Dale, IL). The apparatus was cleaned with 10% ethanol in between animals. The dependent variables were the number of open section entries and total time spent in open sections (defined as the head and front two paws in open sections), which are each indicators of anxiolytic-like behavior in this procedure. A diagram of the zero maze paradigm is included in Fig. 5C.

### Tissue collection

Rats and mice were fasted for 12h before all tissue extractions. Rat brains were rapidly removed from the skull and flash frozen in 30°C isopentane on dry ice, then stored at −80°C to until further processing. Nodose ganglia to be processed for gene expression were flash frozen on dry ice and stored at −80°C to await further processing (see qPCR). Nodose ganglia to be processed for histology were postfixed in ice-cold 4% paraformaldehyde for 2h, transferred to 25% sucrose at 4°C, and remained in sucrose for at least 24h before sectioning (see Histology or FISH). Nodose ganglia for qPCR analyses were flash frozen in −30°C isopentane on dry ice, then stored at −80°C to await further processing.

### Immunoblotting

Tissue punches of brain regions of interest (2.0mm circumference, 1-2mm depth) were collected from brains (knockdown n=13, control n=9) using a Leica CM 1860 cryostat (Wetzlar, Germany) and anatomical landmarks were based on the Swanson rat brain atlas [65]. Tissue punches were enriched with the dorsal cornu ammonis area 1 and dorsal dentate gyrus (dCA1+dDG; Swanson atlas levels 29-30), and the cornu ammonis area 3 (dCA3; atlas levels 29-30). Proteins in brain lysates were separated using sodium dodecyl sulfate polyacrylamide gel electrophoresis, transferred onto poly-vinylidene difluouride membranes, and subjected to immunodetection analysis using enhanced chemiluminescence (Chemidoc XRS, BioRad). A rabbit anti-brain-derived neurotrophic factor antibody (1:500, Santa Cruz Biotechnology, Catalog # sc-20981) was used to evaluate the concentration of brain-derived neurotrophic factor (BDNF) relative to a loading control signal detected by a rabbit anti-β-actin antibody (1:5000, Santa Cruz Biotechnology, Catalog # NB600-503). A rabbit anti-doublecortin antibody (1:500, Abcam, Catalog # ab18723) was used to evaluate the concentration of doublecortin (DCX) relative to a loading control signal detected by an anti-β-tubulin antibody (1:5000, Cell Signaling, Catalog #2128S). Blots were quantified with densitometry analysis using Image J as previously described [17].

### Fluorescent In Situ Hybridization (FISH)

Nodose ganglia (n=3 animals, both right and left ganglia for a total of 6 ganglia) were cut on the Leica CM 1860 cryostat (Wetzlar, Germany) at 20μm and mounted directly onto slides. Slides were washed 5x in KPBS for 5 min each. Sections were then pretreated by incubating them at 37°C in incubation buffer (100mM Tris buffer and 50mM EDTA in distilled deionized water, pH 8) with 0.001% Proteinase K (Sigma P2308) for 30 min, followed by a 3 min wash in incubation buffer alone and a 3 min rinse in 100mM Triethanolamine in water (pH 8). Sections were then incubated with 0.25% acetic anhydride in 100 mM triethanolamine for 10 min at room temperature followed by 2 × 2 min washes in saline-sodium citrate buffer (1% citric acid trisodium/2% sodium chloride in water (pH 7.0)). Slides were dehydrated in increasing concentrations of ethanol solution (50%, 70%, 95%, 100%, 100%) for 3 min each and air-dried prior to hybridization. For hybridization, a hydrophobic barrier was drawn around each section and 3 to 4 drops of the probes (*Ghsr* mRNA probe, *NeuN* mRNA, and *Dapi* mRNA probe, all Advanced Cell Diagnostics; Newark, CA) were placed on each tissue section. Slides were incubated with the probes at 40°C for 3h in a HybEz oven (Advanced Cell Diagnostics). Following a 2 min wash with wash buffer (RNAscope®, Advanced Cell Diagnostics 320058) reagents from RNAscope® Fluorescent Multiplex Detection Reagent Kit (Advanced Cell Diagnostics, 320851) were applied in order to amplify the probe signals, with AMP1 applied for 45 min, AMP2 for 30 min, AMP3 for 45 min, and AMP4 for 30 min. Incubation steps occurred at 40°C, and a 2 min wash followed each amplification step. Slides were coverslipped with ProLong® Gold Antifade mounting medium (Cell Signaling, 9071S). Photomicrographs were acquired using a Nikon 80i (Nikon DS-QI1, 1280×1024 resolution, 1.45 megapixel) under epifluorescent illumination. The percentage of *NeuN*+ nodose ganglia neurons that co-express *Ghsr* was quantified using Nikon Elements software.

### Quantitative polymerase chain reaction (qPCR)

To quantify knockdown of *Ghsr* mRNA expression, quantitative polymerase chain reaction (qPCR) was performed on nodose ganglia, as previously described [66, 67], with left (LNG) and right nodose ganglia (RNG) analyzed separately. Briefly, total RNA for rat nodose ganglia was extracted according to manufacturer’s instructions using the RNeasy Lipid Tissue Mini Kit (Qiagen, Hilden, Germany) for both the shRNA experiments (knockdown n=8 ganglia, control n=12 ganglia, left/right counterbalanced) and the laterality experiments in the rat (n=10 ganglia; left n=5 ganglia, right n=5 ganglia). For the laterality experiments in the mice, total RNA for mouse nodose ganglia (n=8 ganglia; left n=4 ganglia, right n=4 ganglia) was extracted according to manufacturer’s instructions using the RNeasy Micro Kit (Qiagen). RNA was reverse transcribed to cDNA using the Quantitect Reverse Transcription Kit (Qiagen), and amplified using the TaqMan PreAmp Master Mix Kit (ThermoFisher Scientific, Waltham, MA). qPCR was performed with TaqMan Universal PCR Master Mix (Applied Biosystems, Foster City, CA) using the Applied Biosystems QuantStudio 5 Real-Time PCR System (ThermoFisher Scientific). Negative reverse-transcribed samples were generated and all reactions were carried out in triplicate, and control wells without cDNA template were included. The following TaqMan probes were used: Rat *Ghsr:* Rn00821417_m1, Rat *Mchr1*: Rn00755896_m1, Rat *Cnr1*: Rn00562880_m1, Rat *Gapdh*: Rn01775763_g1, Mouse *Ghsr:* Mm00616415_m1, Mouse *Gapdh:* Mm99999915_g1. To determine relative expression values, the 2^-ΔΔ*CT*^ method was used [10], where triplicate Ct values for each sample were averaged and subtracted from those derived from *Gapdh*.

### Spatial foraging task

To test the effect of VAN-specific GHSR knockdown on a food seeking task requiring visuospatial learning and memory, we tested the animals on a novel spatial foraging task modified from the traditional Barnes maze procedure [68]. Briefly, rats are trained on a Barnes maze apparatus (elevated circular maze with 18 holes around the perimeter) with fixed spatial cues on the walls. Rats undergo training (four consecutive training days, two trials per day, trials 120 min apart) to learn the fixed location of a hidden escape tunnel that contains five highly palatable sucrose pellets located in one of the 18 holes around the maze. Rats are tested in a single 2 min probe trial in which the hidden tunnel and sucrose pellets are removed. During the probe trial, normal rats will preferentially investigate the hole in which the sucrose pellets and hidden tunnel were previously located.

### Conditioned place preference

To test the effect of VAN-specific GHSR knockdown on contextual learning for a food reward, rats underwent a conditioned place preference (CPP) behavioral paradigm. Procedures followed the CPP protocol as previously described [69]. Briefly, rats were habituated to the CPP apparatus, which consists of two conjoined plexiglass compartments with a guillotine door in the center (Med Associates, St. Albans, VT). The two compartments (contexts) differ in wall color and floor texture. Habituation occurred with the CPP apparatus door open, allowing them to freely explore each context. Time in each context was measured using AnyMaze Behavior Tracking Software (Stoelting Co., Wood Dale, IL). For each rat, the context that was least preferred during habituation was designated as the food-paired context for subsequent training. CPP training consisted of one 20 min session per day over 16 consecutive days, with eight food-paired and eight non-food paired counterbalanced sessions occurring in total. The CPP apparatus door was closed during all training sessions, restricting rats to one context only. During food-paired training sessions, rats were isolated in the food-paired context with 5 g of palatable food (45% kcal high fat/sucrose diet [D12451, Research Diets, New Brunswick, NJ]) placed on the chamber floor. During non-food paired training sessions, rats were isolated in the non-food-paired context with no food present. The CPP test occurred two days after the last training session using a between-subjects design. During testing, the CPP apparatus door was opened, and the rats were allowed to freely explore both contexts during a 15 min trial. For all habituation, training and testing, the apparatus was cleaned with 10% ethanol in between animals. Time in each context was measured using AnyMaze Behavior Tracking Software (Stoelting Co., Wood Dale, IL). The dependent variable was the percentage shift in preference for the food-associated context during testing compared with the baseline session.

### Statistical analyses

Data are presented as mean ± SEM. For comparisons of over time of body weight, food intake (cumulative 24h intake, meal pattern, meal entrainment), gastric emptying, blood glucose, and postprandial insulin, groups were compared using a two-way repeated measures ANOVA (group x time) and significant ANOVAs were analyzed with a Fisher’s LSD posthoc test where appropriate. For comparisons of food intake after IP ghrelin treatment, groups were compared using a two-way repeated measures ANOVA (surgery x treatment). Paired two-sample two-tailed Student’s t-tests were used for comparison of within-animal neural responses in the DAPR experiments. Unpaired, two-sample, two-tailed Student’s t-tests were used for all other analyses Differences were considered statistically significant when p<0.05. Outliers were identified as being more extreme than the Median +/- 1.5 * Interquartile Range. Assumptions of normality, homogeneity of variance, and independence were met when required.

---

**Figure S1.**
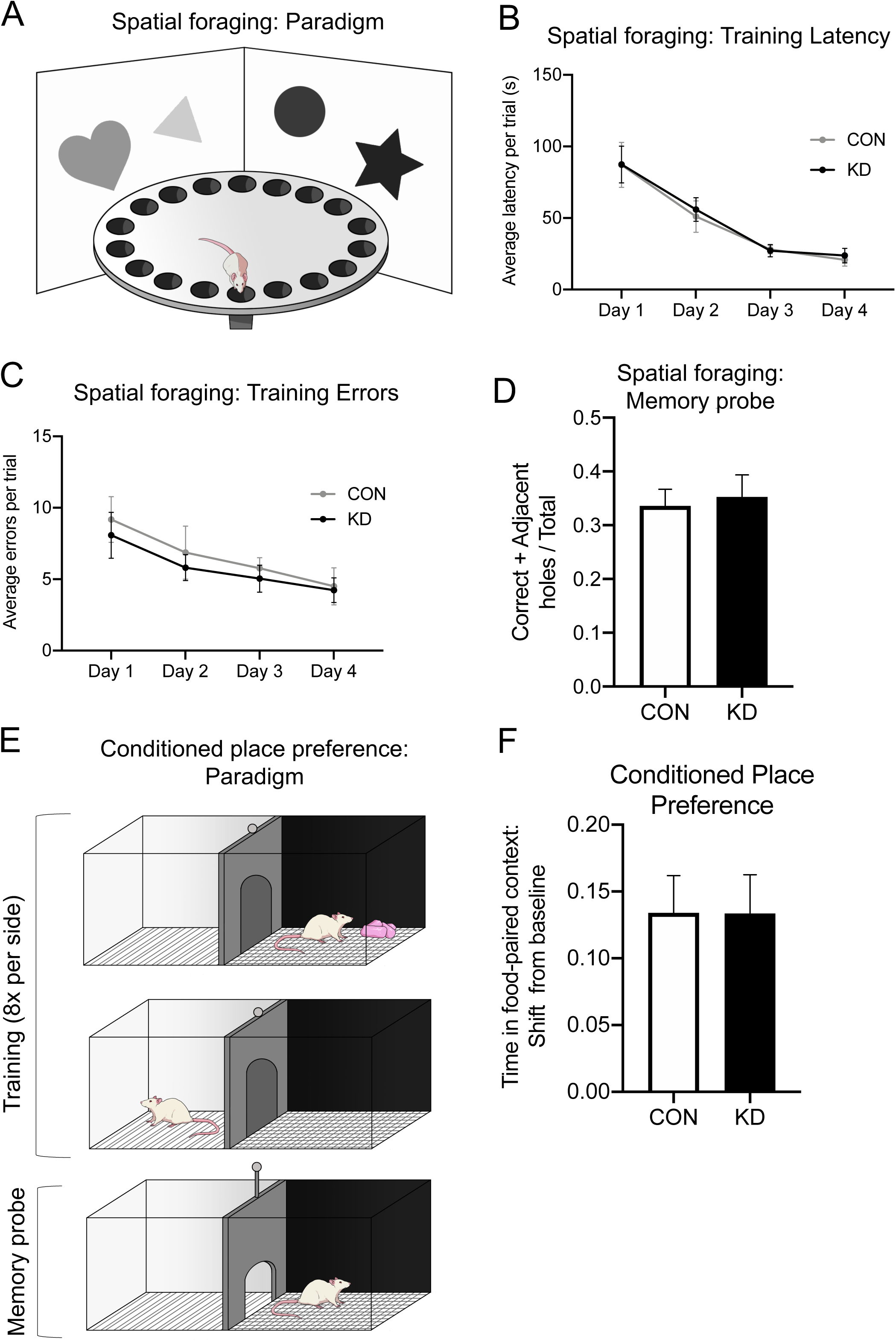
VAN-specific GHSR knockdown does not affect appetitive learning and memory for palatable food. In a novel spatial foraging task (A), there were no group differences in learning, as measured by latency to correct hole (B) and errors before correct hole during training (C). There were also no group differences in memory, as measured by the correct + adjacent holes investigated / total holes investigated during a 2 min memory probe (D). In the conditioned place preference task for high fat diet (E), results showed no differences between VAN-specific GHSR knockdown and controls in preference for food-paired context (time spent in context, shift from baseline) (F). All data presented as mean +/- SEM.

**Figure S2.**
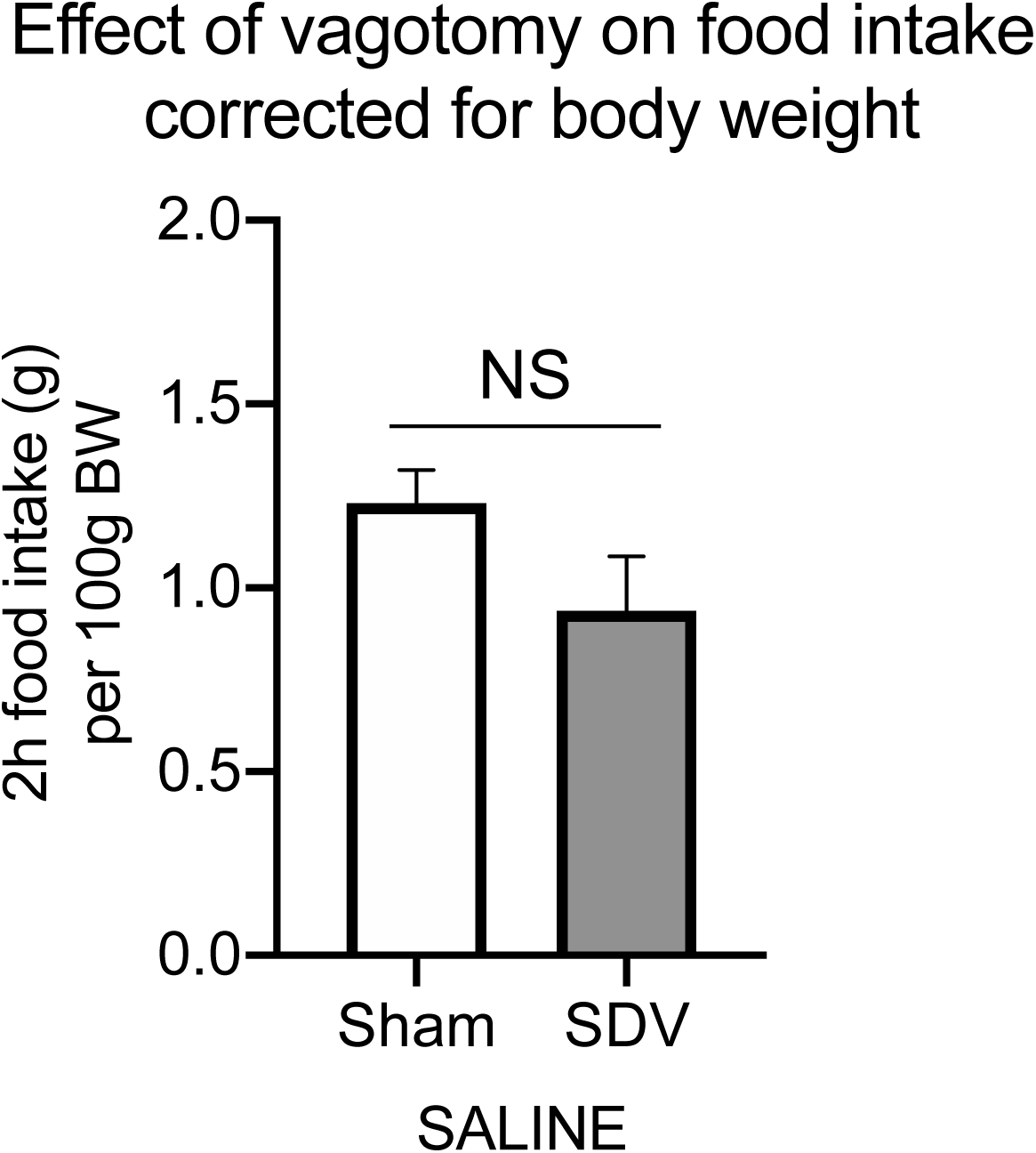
There is no difference between 2h dark cycle food intake after saline injection (control) between sham and vagotomized animals when corrected for body weight (A). Data presented as mean +/- SEM.

**Figure S3.**
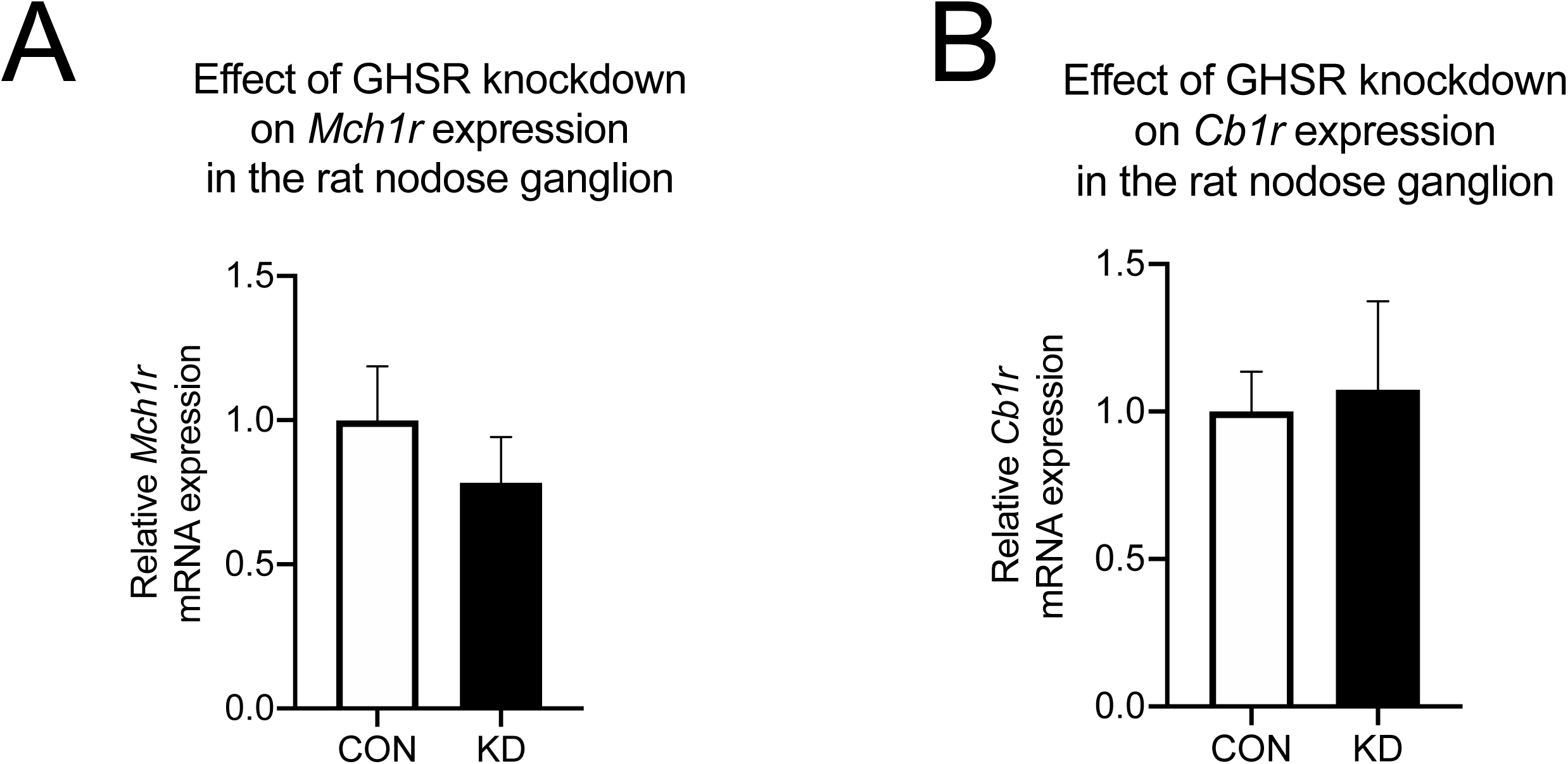
VAN-specific GHSR knockdown does not affect expression of *Mch1r* (A) or *Cb1r* (B). All data presented as mean +/- SEM.

